# Universality and non-universality of the growth law

**DOI:** 10.1101/2021.12.02.471021

**Authors:** Qirun Wang, Jie Lin

**Affiliations:** Center for Quantitative Biology, Peking University, Beijing, China; Peking-Tsinghua Center for Life Sciences, Peking University, Beijing, China

## Abstract

An approximately linear relationship between the fraction of ribosomal proteins in the proteome (*ϕ_R_*) and the growth rate (*μ*) holds in proliferating cells when the nutrient quality changes, often referred to as a growth law. While a simple model assuming a constant translation speed of ribosomes without protein degradation can rationalize this growth law, real protein synthesis processes are more complex. This work proposes a general theoretical framework of protein synthesis, taking account of heterogeneous translation speeds among proteins and finite protein degradation. We introduce ribosome allocations as the fraction of active ribosomes producing certain proteins, with two correlation coefficients respectively quantifying the correlation between translation speeds and ribosome allocations, and between protein degradation rates and mass fractions. We prove that the growth law curve generally follows *ϕ_R_* = (*μ* + *c*_1_)/(*c*_2_*μ* + *c*_3_) where *c*_1_, *c*_2_, and *c*_3_ are constants depending on the above correlation coefficients and the translation speed of ribosomal proteins. Our theoretical predictions of *ϕ_R_* agree with existing data of *Saccharomyces cerevisiae*. We demonstrate that when different environments share similar correlation coefficients, the growth law curve is universal and up-bent relative to a linear line in slow-growth conditions, which appears valid for *Escherichia coli*. However, the growth law curve is non-universal and environmental-specific when the environments have significantly different correlation coefficients. Our theories allow us to estimate the translation speeds of ribosomal and non-ribosomal proteins based on the experimental growth law curves.

---

Cells can adapt to different environments and alter the expression levels of multiple genes correspondingly. The genome-wide gene expression profile can change significantly as cells switch between different environments. However, proliferating cells, including bacteria and unicellular eukaryotes, exhibit a simple growth law as the nutrient quality changes: an approximately linear relation exists between the fraction of ribosomal proteins in the proteome (*ϕ_R_*) and the growth rates (*μ*), *ϕ_R_* = *μ*/*κ* + *ϕ*_0_ [1–6]. This growth law can be rationalized by a simple translation model (STM): ribosomes are engaged in translation with a constant translation speed that is proportional to *κ* [2, 4]. *ϕ*_0_ represents the fraction of inactive ribosomes that are not producing proteins, independent of environments in the STM. While the STM is simple and intuitive, it appears to break down in slow-growth conditions of *Escherichia coli* (doubling time longer than 60 mins at 37°C) in which more ribosomes are produced than the expectation from the STM [7].

We note that there are two important biological features (if not all) beyond the STM, which, as we show in this work, are crucial to interpret the experimental data of *ϕ_R_* versus *μ* (the growth law curve). The first is the heterogeneous translation speeds of ribosomes producing different proteins. Recent studies demonstrated that the translation speeds are highly heterogeneous among different proteins due to multiple mechanisms, including codon usages [8] and amino acid compositions [9]. Because of the universalities of these mechanisms, one expects that heterogeneous translation speeds among proteins are universal across different organisms. In particular, the translation speeds of ribosomal proteins are significantly slower than the average translation speed over non-ribosomal proteins due to the abundance of positively charged amino acids on ribosomal proteins [9]. Nowadays, the ribosome profiling technique allows us to quantify the allocation of ribosomes towards the production of different proteins. These experimental techniques enable us to rethink the growth law in the presence of heterogeneity in translation speeds [9].

The second feature is finite protein degradation rates. The STM neglects protein degradation and predicts that at zero growth rate, *ϕ_R_* = *ϕ*_0_ so that all ribosomes are inactive. However, this contradicts with experiments of nongrowing bacteria in which significant translation activities are observed [10]. Protein degradation must be considered at zero growth rate to balance protein production to ensure a constant protein mass. Therefore, protein degradation should be important to the growth law, at least in slow-growth conditions.

In this work, we show that the heterogeneous translation speeds and finite protein degradations significantly influence the growth law connecting the fraction of ribosomal proteins and the growth rate when the nutrient quality changes. The fractions of ribosomal proteins *ϕ_R_* are generally different in different environments, even if they lead to the same growth rates. Besides the growth rate, *ϕ_R_* depends on two correlation coefficients among proteins. One is between the translation speeds and ribosome allocations, and the other is between the correlation coefficient between protein degradation rates and mass fractions. We compute the above correlation coefficients using proteomics and ribosomal profiling datasets of *S. cerevisiae* [11]. Interestingly, we find that the correlation between the translation speed and ribosome allocations become stronger when the growth rate decreases; namely, cells tend to produce more proteins with higher translation speeds in poor nutrient. In contrast, the correlation between the protein degradation rates and mass fractions is almost independent of growth rates.

We derive the general form of growth law involving the above correlations. We demonstrate that for environments with similar correlation coefficients, the growth law curve is universal and has the following form, *ϕ_R_* = (*μ* + *c*_1_)/(*c*_2_*μ* + *c*_3_) where *c*_1_, *c*_2_, and *c*_3_ are constants depending on the above correlation coefficients and the translation speed of ribosomal proteins. We prove that the growth law curve must be monotonically increasing and convex, which justifies the upward bending of the growth law curve of *E. coli* observed in slow-growth conditions relative to a linear line [7]. However, if the experiments are implemented in multiple environments with dramatically different correlation coefficients, the growth law curve is generally non-universal and environmental-specific. Our analysis of experimental data suggests that this scenario may apply to *S. cerevisiae*. Our theories allow us to fit the experimentally measured growth law curves by our model predictions, from which we can estimate the translation speed of ribosomal and non-ribosomal proteins. Consistent with direct experimental measurements [9], the estimated translation speed of ribosomal proteins is indeed much slower than non-ribosomal proteins.

## RESULTS

### Model of protein synthesis

Given a constant environment, we consider a population of cells with a constant growth rate, and the protein synthesis processes are in a steady state. Ribosome profiling allows us to quantify the fraction of ribosomes in the pool of total active ribosomes producing protein *i*, which we call ribosome allocation *χ_i_*. Here the index *i* represents one particular protein *i*. Mass spectrometry also allows us to measure the mass fractions *ϕ_i_* of all proteins in the proteome [12]. The elongation rate of ribosomes on the corresponding mRNAs is *v_i_*, which is the number of translated amino acids per unit time. Note that *v_i_* is the averaged elongation rate over the sequence of the corresponding mRNA so that each protein has one *v_i_*. We also assume that protein *i* degrades with a constant rate *α_i_*. The mass production rate of protein *i* becomes

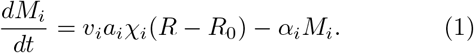

Here *R* is the number of ribosomes, and *R*_0_ is the number of inactive ribosomes. *α_i_* is the averaged mass of amino acids over the sequence of protein *i*. In the following analysis, we define *k_i_* = *v_i_ α_i_* as the amino acid mass-weighted translation speed and denote it as the translation speed for simplicity. Our model is summarized in Figure 1.

**FIG. 1.**
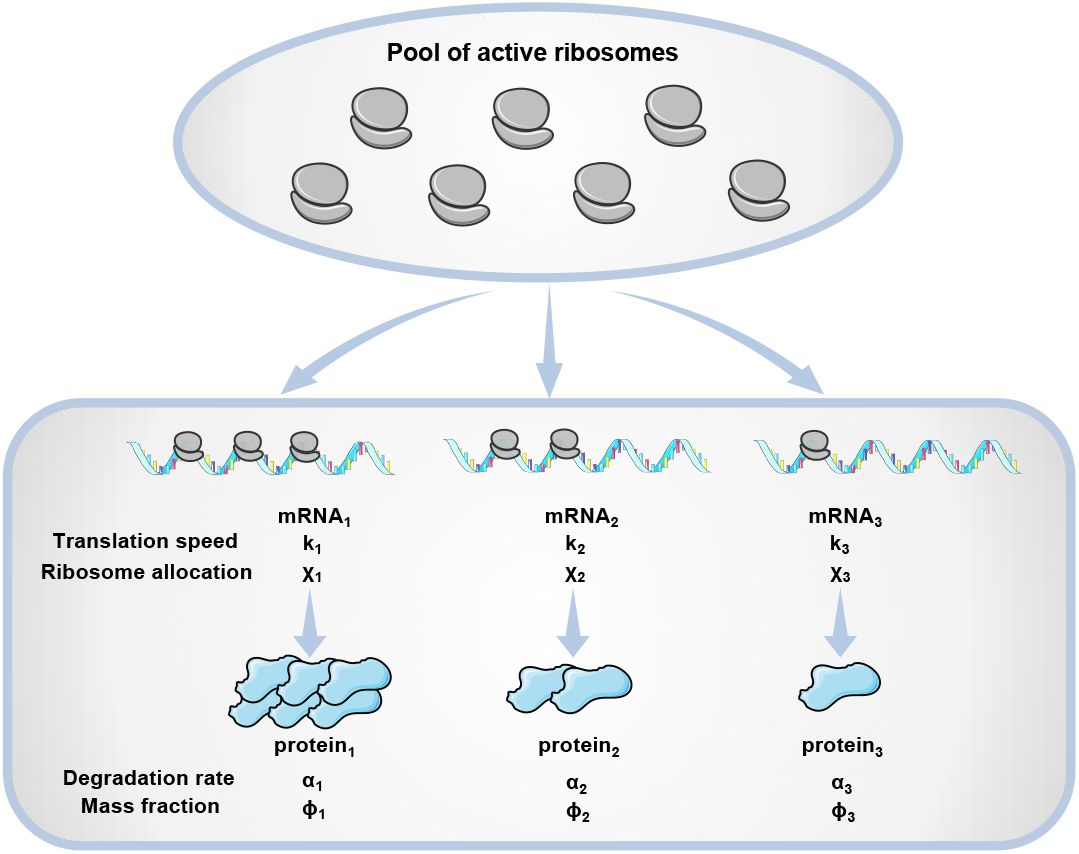
Given a constant environment, cells actively allocate different fractions of active ribosomes (*χ_i_*) to translate mRNAs corresponding to different proteins. In general, the translation speeds *k_i_* are heterogeneous among proteins. *α_i_* is the degradation rate of protein *i. χ_i_, k_i_* and together determine the mass fraction of protein *i*. The ribosome allocation strategies reflect the adaption of cells to different environments. In this schematic, we show three proteins for simplicity.

Recently, Dai et al. showed that for *E. coli* the translation speeds of many proteins decrease as the growth rate decreases, but maintain finite values at zero growth rate [7]. They proposed a model in which the translation speeds are the same for all proteins and depend on the ribosomal fraction *ϕ_R_* in a Michaelis-Menten way, consistent with their experimental data. However, their model predicts a downward bending of the growth law curve in slow-growth conditions relative to a linear line, in contrast to the upward bending observed experimentally. To reconcile the conflict, they proposed that the fraction of inactive ribosomes *ϕ*_0_ increases as the growth rate decreases, generating the upward bending of the growth law curve. However, as far as we know, there is no direct experimental evidence supporting a larger fraction of inactive ribosomes *ϕ*_0_ in slow-growth conditions than in fast-growth conditions. Interestingly, no noticeable bending is observed in the growth law curve of *S. cerevisiae* [6], suggesting that the upward bending of the growth law curve in slow-growth conditions may not be universal across organisms, consistent with our theoretical predictions as we show later.

We remark that a growth-rate dependent translation speed is undoubtedly a mechanism that the STM breaks down. However, in this work, we focus on the effects of heterogeneous translation speeds *k_i_* and finite degradation rates *α_i_*. Therefore, we assume them to be invariant of environments. We also mainly consider the effects of nutrient quality and do not consider the impact of antibiotics in this work, which can decrease the overall effective translation speed and increase *ϕ_R_* as the growth rate decreases [4]. Thanks to the simplicity of our protein synthesis model, it can be analytically solved, and the predictions are intriguing as we show later.

We define the total protein mass *M* = Σ_*i*_ *M_i_*, and the protein mass fraction *ϕ_i_* = *M_i_*/*M*. Using Eq. (1), we find the values of *ϕ_i_* in the steady state as (see detailed derivations in Appendix A)

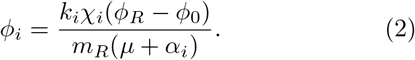

Here *μ* is the growth rate of the total protein mass 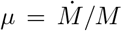, and *m_R_* is the total amino acid mass of a single ribosome. Since all proteins grow in the same rate in the steady-state, the growth rates of protein *i* defined as 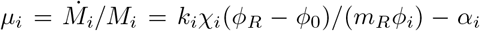 must be equal to *μ*, which can be easily verified using Eq. (2). In the following, *i* = 1 is reserved for ribosomal proteins so that *ϕ*_1_ = *ϕ_R_* and *μ_R_* = *μ_R_* = *k_R_χ_R_*(1-*ϕ*_0_/*ϕ_R_*)/*m_R_*–*α_R_*. Here, *k_R_* and *α_R_* are the effective translation speed, and degradation rate of the coarse-grained ribosomal protein averaged over all ribosomal proteins. They are approximately independent of environments due to the tight regulation of relative doses of different ribosomal proteins [13] and their generally low degradation rates.

Given the ribosome allocations *χ_i_*, the protein degradation rates *α_i_* and the translation speeds *k_i_*, one obtains a unique solution of *ϕ_i_* and *μ*. We can express the growth rate as *μ* = Σ_*i*_ *ϕ_i_μ_i_* and rewrite Eq. (2) to obtain the expression of *ϕ_R_* as (see detailed derivations in Appendix B)

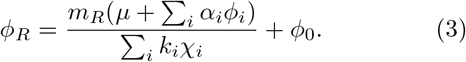

Here, *ϕ*_0_ is the mass fraction of inactive ribosomes, which we assume to be constant in the following. It is easy to find that if all proteins have the same translation speed (*k_i_* = *k* for all *i*) and protein degradations are negligible (*α_i_* = 0), Eq. (3) is reduced to the STM result.

### Effects of heterogeneous translation speeds

To better understand the effects of heterogeneous translation speeds and degradation rates, we choose to study them separately. Therefore, we first simplify the model by taking *α_i_* = 0 for all proteins and only consider the effects of heterogeneous translation speeds *k_i_*. We rewrite 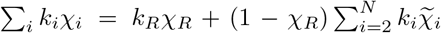 in Eq. (3). Here, *N* is the number of genes and 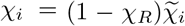 so that 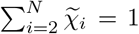. *k_R_* is the translation speed of ribosomal proteins. In the following, we define 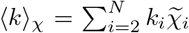 as the *χ*-weighted average translation speed over all non-ribosomal proteins. As we derive in Appendix C, the fraction of ribosomal proteins can be written exactly as a Hill function of the growth rate:

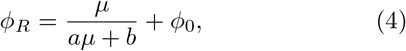

where

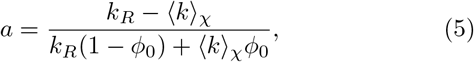

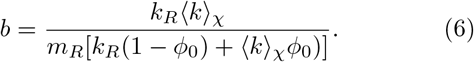

We are particularly interested in the sign of *a* because it determines the shape of the *ϕ_R_*(*μ*) curve. If *k_R_* is smaller than 〈*k*〉_χ_, *a* is negative so that the second derivative of the *ϕ_R_*(*μ*) curve is positive. In other words, the *ϕ_R_*(*μ*) curve is upward bent in slow-growth conditions.

〈*k*〉_*χ*_ depends on both the elongation speeds *k_i_* and the ribosome allocations *χ_i_*. To find its value, we further rewrite 〈*k*)_*χ*_ = 〈*k*〉(1 + *I_χ,k_*). Here 〈*k*〉 is the arithmetic average of translation speeds over all non-ribosomal proteins, which is constant and independent of environments. *I_χ,k_* is a metric we use to quantify the correlation between the ribosome allocations and the translation speeds:

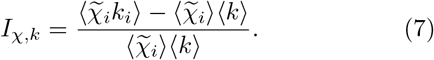

Here, the bracket represents an average over all non-ribosomal proteins. Because the ribosomal allocations *χ_i_* are generally different in different environments, we use *I_χ,k_* to characterize an environment. Imagine that we grow cells in multiple environments with equal *I_χ,k_*. We find that as long as *I_χ,k_* is not too close to −1, which we confirm later using experimental data, *a* is always negative since the translation speed of ribosomal proteins *k_R_* is much lower than 〈*k*〉 [9]. Therefore, Eq. (4) predicts an upward bending of the *ϕ_R_*(*μ*) curve in slow-growth conditions.

We verify the above theoretical predictions by numerically simulating the model of protein synthesis (Appendix E). The translation speeds are randomly sampled among proteins and fixed for all environments, with *k_R_* < 〈*k*〉. We randomly sample *χ_i_* for each environment and compute the resulting growth rate *μ* and protein mass fractions *ϕ_i_*. We show the results from environments with preselected *I_χ,k_*, which agree well with the theoretical formula Eq. (4) (Figure 2a).

**FIG. 2.**
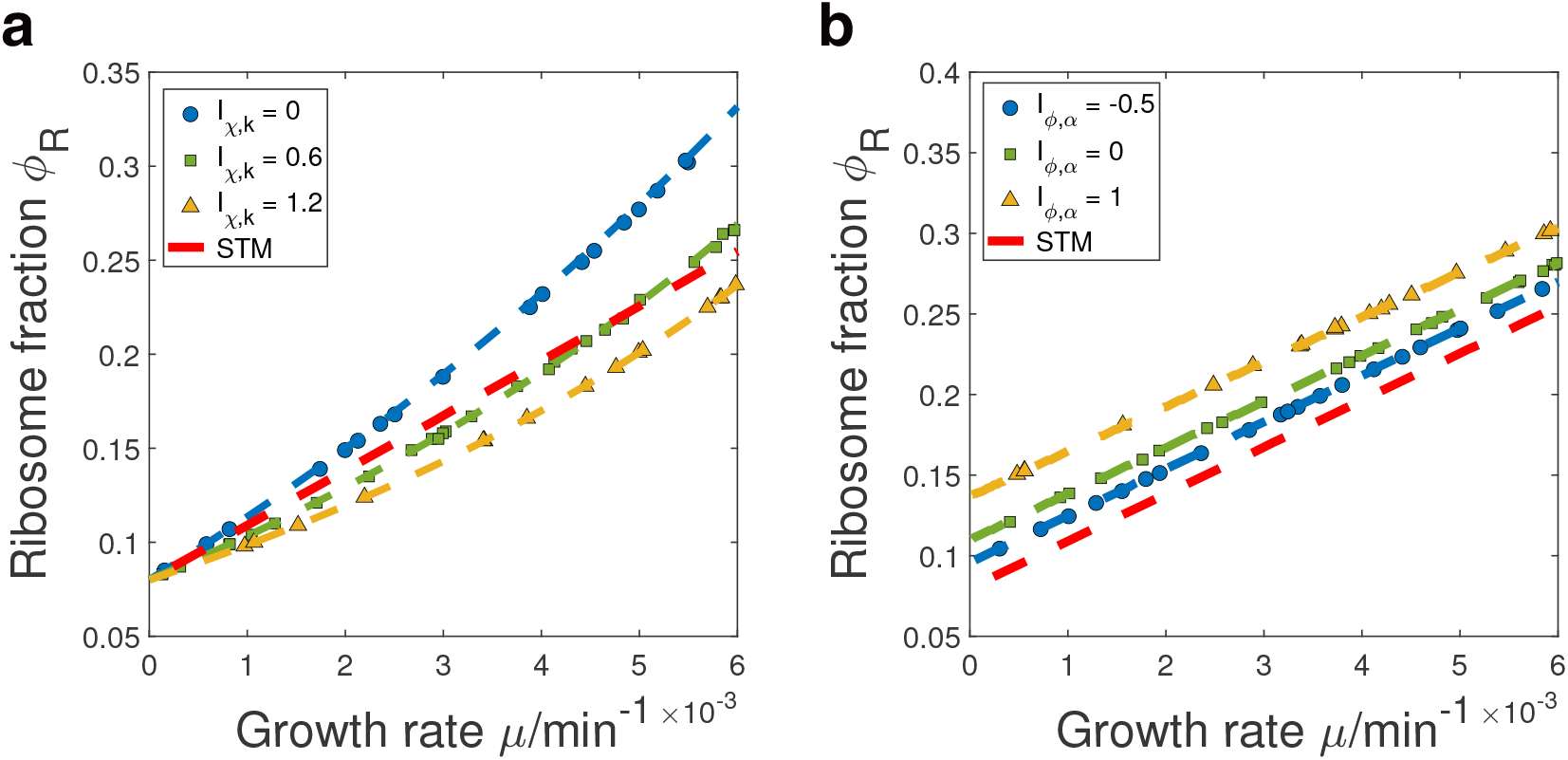
Numerical simulations of the growth law curves. (a) We simulate the case of heterogeneous translation speeds and compare our numerical simulations with model predictions (dashed lines). Each data point has its own randomly sampled and we show the results with preselected *I_χ,k_* values. The red dash line represents the predictions of the STM in which all proteins have the same translation speed 〉*k*〈. (b) Same analysis in which we simulate the case of finite protein degradation rates.

### Effects of finite protein degradation rates

We now discuss the effects of finite protein degradation rates. For simplicity, we assume that the translation speeds are homogeneous and equal to *k* for all proteins. We rewrite the Σ_*i*_ *α_i_ϕ_i_* term in Eq. (3) such that 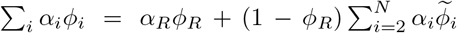. Here, 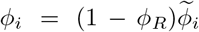 so that 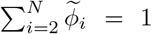. We define the *ϕ*−averaged degradation rates over all non-ribosomal proteins as 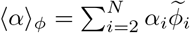. Therefore, Eq. (3) can be written as

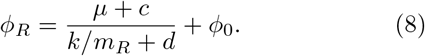

where

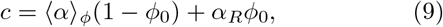

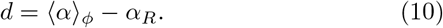

To find the sign of *d*, we further rewrite 〈*α*〉_*ϕ*_ as 〈*α*〉_*ϕ*_ = 〈*α*〉(1+*I_ϕ,α_*) where 〈*α*〉 is the arithmetic average of degradation rates over all non-ribosomal proteins. *I_ϕ,α_* is a metric we use to characterize an environment by quantifying the correlation between the protein mass fractions and degradation rates:

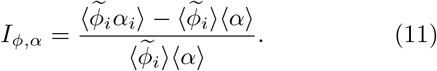

Here, the bracket represents an average over all non-ribosomal proteins.

Imagine that we grow cells in multiple environments with equal *I_ϕ,α_*. We assume that the degradation rate of ribosomal protein *α_R_* is slower than the average of non-ribosomal proteins 〈*α*〉, which is biologically reasonable since ribosomal proteins are generally non-degraded. Therefore, as long as *I_ϕ,α_* is not too close to −1, which we confirm later using experimental data, d is positive since *α_R_* is always smaller than 〈*α*〉_*ϕ*_. Therefore, our model predicts that the growth law curve is linear given a constant *I_ϕ,α_* and finite protein degradation decreases the slope relative to the STM. The intercept at *μ* = 0 is also larger than *ϕ*_0_. Therefore, a finite fraction of ribosomes are still actively translating at zero growth rate. We verify the above theoretical predictions by numerically simulations and randomly sample the protein degradation rates that are fixed for all environments, with *α_R_* < 〈*α*〉 satisfied. We show the results from environments with preselected *I_ϕ,α_* and our theoretical predictions Eq. (8) are nicely confirmed (Figure 2b).

### The full model

We now consider the full model with both the heterogeneities in the translation speeds and protein degradation rates. We find that the growth law curve has the following general form,

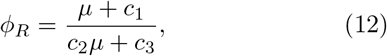

where the expression of the constants, *c*_1_, *c*_2_ and *c*_3_ are shown in Appendix D. We prove that given fixed *I_χ,k_* and *I_ϕ,α_* (as long as they are not too close to −1), the growth law curve must be monotonically increasing and convex, which suggests an upward bending in slow-growth conditions (Appendix D). The simulation results again match well with the theoretical predictions (Figure 3a).

**FIG. 3.**
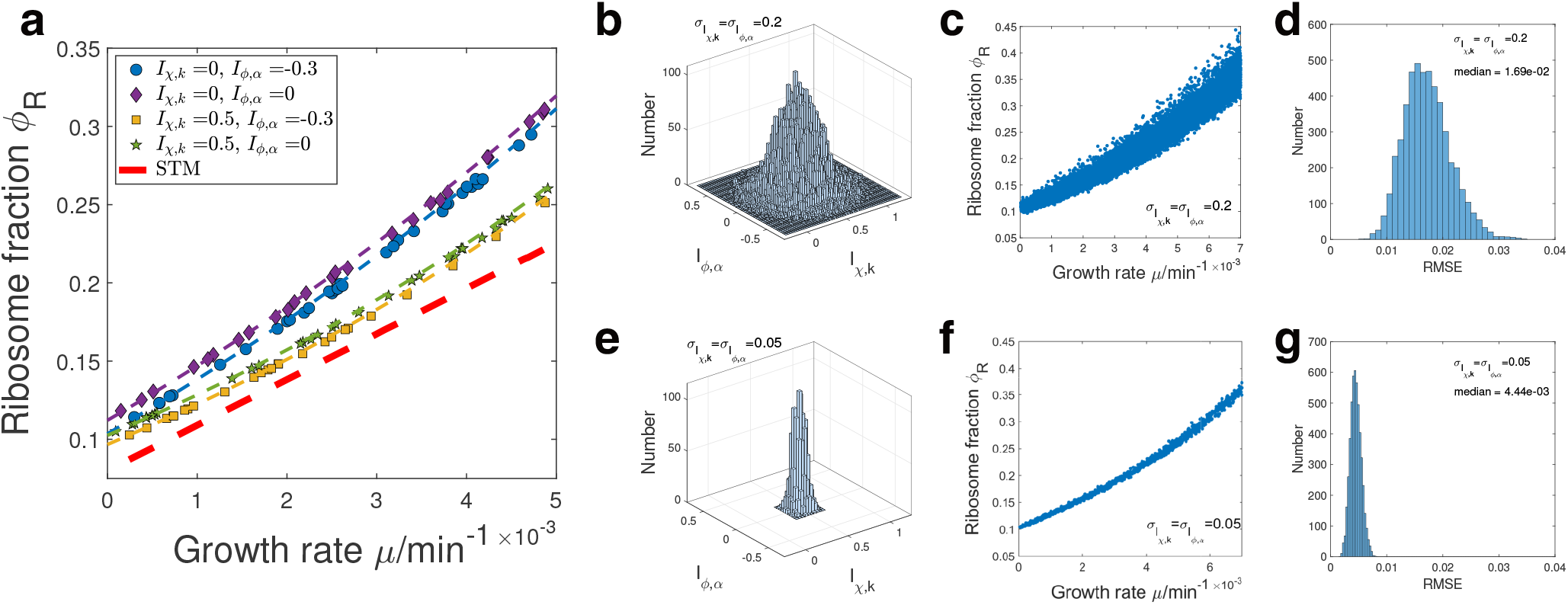
Numerical simulations of the growth law curves with both heterogeneous translation speeds and protein degradation rates. (a) Numerical simulations with preselected *I_χ,k_* and *I_ϕ,α_*. The red dashed line is the prediction of the STM and other dashed lines represent our model predictions. (b) and (e) Two-dimensional Gaussian distribution of randomly sampled *I_χ,k_* and *I_ϕ,α_*. The mean of *I_χ,k_* is 0.5 and the mean of *I_ϕ,α_* is 0. The standard deviations *σ* are indicated in the legends. (c) and (f) The resulting growth law curve where each point has randomly sampled *I_χ,k_* and *I_ϕ,α_* from (b) and (e). (d) and (g) The distributions of the fitting RMSE corresponding to randomly chosen points in (c) and (f).

In real situations, we remark that the actual growth curve shape depends on the particular environments. To verify this, we compute the resulting growth law curve with multiple environments, and the *I_χ,k_* and *I_ϕα_* of each environment are randomly sampled from Gaussian distributions (Figure 3b and e) (Appendix E). We find that when the Gaussian distributions have large standard deviations, the growth law curve is non-universal and depends on the particular chosen environments (Figure 3c). This means that if we randomly pick some environments from Figure 3c, the resulting growth law curves are generally different. In contrast, when the Gaussian distributions have small standard deviations, the growth law curve is well captured by our theoretical predictions Eq. (12), because the environments share similar *I_χ,k_* and *I_ϕ,α_* (Figure 3f).

To quantify the effects of heterogeneous *I_χ,k_* and *I_ϕ,α_* across environments, we repeatedly sample 20 random points from Figure 3c, f and fit them using Eq.(12) (Appendix E). We find that when the chosen environments have significantly different *I_χ,k_* and *I_ϕ,α_*, the median root mean squared error RMSE = 1.69 × 10^-2^ (Figure 3d). In contrast, in the case of similar environments, RMSE = 4.44 × 10^-3^ (Figure 3g). The above results suggest that we can use the fitting error as a criterion of the universality of the growth law curve, which we apply to the experimental data later.

### Experimental tests of theories

In this section, we test our model using published datasets of *S. cerevisiae* [14] (Appendix F). For each strain and nutrient quality, we computed the correlation coefficients between the translation speeds and ribosome allocations *I_χ,k_*, and the correlation coefficients between the protein degradation rates and protein mass fractions *I_ϕ,α_*. Given the values of *μ*, *I_χ,k_*, and *I_ϕ,α_*, we predicted the fraction of ribosomal proteins *ϕ_R_* using Eq. (12) (Figure 4a and e). We note that there is one parameter *ϕ*_0_ that is not known experimentally. Interestingly, by choosing a common *ϕ*_0_ = 0.048, our model predictions nicely match the experimental measured values of *ϕ_R_* (with one data point slightly above the theoretical prediction). We find that regardless of the data processing procedures, the relative relationships between the predicted curves always agree with that of the experimental values (Appendix F and Supplementary Figure S1).

**FIG. 4.**
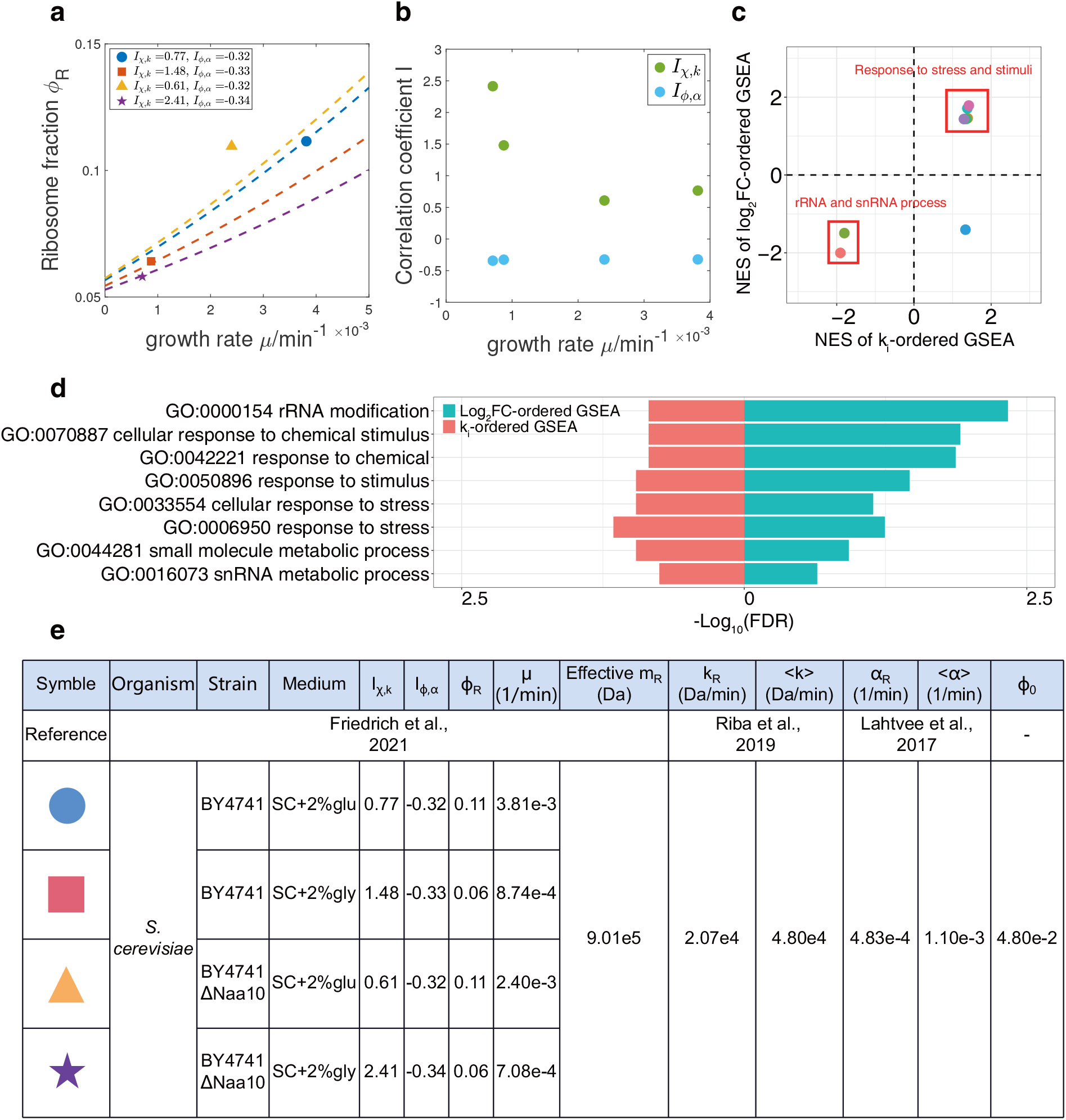
Experimental analysis and theoretical predictions. (a) Experimental measured *ϕ_R_* of *S. cerevisiae* along with the predictions (dashed lines) of our model. (b) The growth rate dependence of the correlation coefficients *I_χ,k_* and *I_ϕα_*. (c) The normalized enrichment score (NES) of GSEA of enriched gene sets. A positive NES of *k_i_*-ordered GSEA means that the genes in the corresponding gene set are enriched in the regime of higher *k_i_*. A positive NES of log_2_FC-ordered GSEA means that the genes in the corresponding gene set are enriched in the regime of increasing *χ_i_* when the nutrient changes from glucose to glycerol. (d) The enriched gene sets with their false discovery rate (FDR) q values of the single-sided permutation test. The higher the – log_10_(FDR) value is, the more likely a gene set is enriched. (e) Summary of the multiple computed variables and parameters in the analysis of experimental data. Note that the effective mass of ribosomal proteins *m_R_* is calculated based on molecular weights of ribosomal proteins detected in the proteome (Appendix F). SC, synthetic complete medium. Glu, glucose. Gly, glycerol.

Our model is simplified as we assume that the translation speeds and protein degradation rates do not depend on environments. Remarkably, our model predictions still quantitatively match the experimental observations, suggesting that our assumptions may be good approximations for most situations. While our model cannot predict the growth rate dependence of *ϕ*_0_, our results show that a constant fraction of inactive ribosomes is consistent with existing datasets of *S. cerevisiae*.

Interestingly, we found that *I_ϕ,α_* ≈ −0.33 for all the conditions we computed. However, *I_χ,k_* are negatively correlated with the growth rates, suggesting cells tend to allocate more ribosomes to translate mRNAs with higher *k_i_* in poor nutrient conditions (Figure 4b). To find out what genes acquire more resources when the environment is shifted, we perform Gene Set Enrichment Analysis (GSEA) [15, 16] for wide type cells (Appendix F) and find that 8 gene sets from the Gene ontology (GO) [17, 18] database are enriched in both the GSEA where genes are ordered by *k_i_* (denoted as *k_i_*-ordered GSEA) and the GSEA where genes are ordered by log_2_ fold change (log_2_FC) of *χ_i_* (denoted as log_2_FC-ordered GSEA) (Figure 4d).

We find that five gene sets related to stress response are enriched in the regime of higher *k_i_* and increasing *χ_i_* when the environment is changed from 2% glucose to 2% glycerol (Figure 4c). This is consistent with the environmental stress response (ESR) of *S. cerevisiae* as an adaptation to the shifts of environments [19]. We propose that higher translation speeds of stress response genes enable cells to respond rapidly to environmental changes, which is evolutionarily advantageous. We also find two gene sets related to the rRNA process are enriched in the regime of lower *k_i_* and decreasing *χ_i_* (Figure 4c). This is consistent with the lower *ϕ_R_* in slow-growth conditions (Figure 4a). We also perform GSEA for ΔNaa1O cells and get similar results (Supplementary Figure S2).

### Applications of theories

An important application of our theories is that one can estimate the translation speeds by fitting the experimental growth law curve to our model prediction Eq. (12) (Appendix G). Because there are 6 unknown parameters in the definition of *c*_1_, *c*_2_, and *c*_3_ (Eq. (23-25)), we can estimate 3 of the parameters given the values of the other 3. For the *S. cerevisiae* data from Ref. [6], we use the experimentally measured degradation rate of ribosomal proteins *α_R_* and the mass of ribosomal proteins *mR* as given. We approximate the *ϕ*-averaged degradation rate 〈*α*〉_*ϕ*_ by 〈*α*〉(1 + *I_ϕ,α_*) where *I_ϕ,α_* = −0.33, and this is justified by the observations that *I_ϕ,α_* is largely independent of environments (Figure 4a). We find that the fitted parameters *c*_1_, *c*_2_ and *c*_3_ having a wide range of 95% confidence intervals (Figure 5a) with RMSE =1.35 × 10^-2^, which suggests that the growth law curve is non-universal according to our simulations (Figure 3d). Indeed, the inferred values of *ϕ*_0_, *k_R_* and 〈*k*〉_*χ*_ have very large error bars (Figure 5c). We also just fit the C-limiting data points in Figure 5a [6] and obtain similar results (Supplementary Figure S3).

**FIG. 5.**
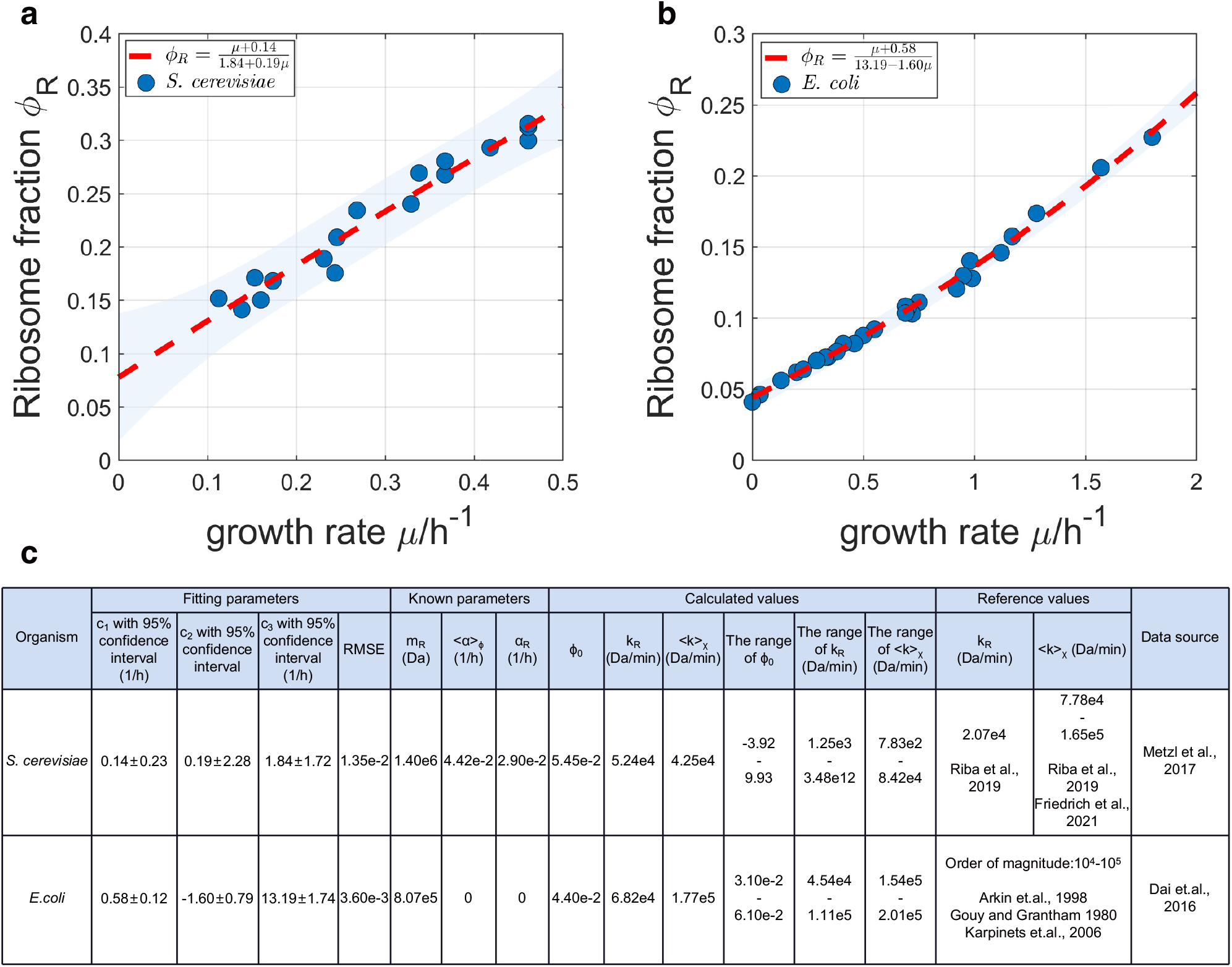
The full model fits different datasets. (a) The non-linear fitting to data from Ref. [6]. The shadow represents the 95% prediction interval. (b) The non-linear fitting to data from Ref. [7]. The shadow is the same as in (a). (c) Detailed fitting results of (a) and (b). Note that the reference value of 〈*k*〉_*χ*_ of (a) is approximated by 〈*k*〉(1 + *I_χ,k_*) where the range of *I_χ,k_* can be found in Figure 4c.

We also apply our theories to *E. coli* [7] (Figure 5b). Because most proteins are non-degradable in bacteria [20, 21], we set *α_R_* and 〈*α*〉_*ϕ*_ as 0, and the mass of ribosomal protein *mR* = 8.07 × 10^5^ *Da* [12]. In this case, the fitted parameters have much smaller range of 95% confidence intervals with RMSE = 3.60× 10^-3^. The estimated *k_R_*, and 〈*k*〉 are consistent with previous studies [22–24] (Figure 5c). Our analysis of experimental data demonstrates that the translation speed of ribosomal proteins is indeed smaller than the *χ*-averaged translation speed, in agreement with experimental observations [9]. Our results suggest that *E. coli* has similar values of *I_χ,k_* and *I_ϕ,α_* in the chosen environments of Ref. [7] so that it has a universal growth law curve. In contrast, *S. cerevisiae* appears to have significantly different *I_χ,k_* and *I_ϕ,α_* across different environments of Ref. [6] so that the growth law curve depends on the chosen environments and therefore non-universal.

## Discussion

In this work, we go beyond the simple translation model and take account of the heterogeneous translation speeds and finite protein degradation. Given the translation speeds and protein degradation rates, our model is completely general and virtually applies to any cells, including both proliferating cells (*μ* > 0) and nonproliferating cells (*μ* = 0). In this work, we mainly consider the scenario in which the growth rate changes due to the nutrient quality and the fraction of ribosomal proteins (*ϕ_R_*) increases monotonically as the growth rate increases.

We demonstrate that the growth law curve is, in general, nonlinear and has the form Eq. (12). In particular, the main effect of heterogeneous translation speeds is making the growth law curve up-bent relative to the STM. The main effect of protein degradation is reducing the slope and increasing the intercept relative to the STM. The actual shape of the growth law curve depends on two correlation coeffcients: one is between the ribo-some allocations and the translation speeds (*I_χ,k_*); the degradation rates (*I_ϕ,α_*)o By analyzing the dataset from [14], we found that *I_ϕ,α_* is independent of growth rate, while *I_χ,k_* appears to be negatively correlated with the growth rate. This means that cells tend to produce proteins with faster translation speeds in slow-growth conditions, which can be an economic strategy and under evolutionary selection. Remarkably, our theoretical pre-dictions of ϕR can reasonably match the experimentally measured values [14], with a common fraction of inactive ribosomes *ϕ*_0_. Our results imply that the fraction of inactive ribosomes may be constant across different nutrient qualities.

We apply our model predictions to the growth law curves of *S. cerevisiae* [6] and *E. coli* [7]. In the former case, the fitting of data to our model prediction is subject to significant uncertainty. This agrees with the computed *I_χ,k_* that are variable across conditions using the ribosome profiling and mass spectrometry data from [14]. In contrast, the fitting of *E. coli* data exhibits a much smaller uncertainty, suggesting that common *I_χ,k_* and *I_ϕ,α_* may apply to all the nutrient qualities used in the experiments of Ref. [7]. This is to be tested when genome-wide measurements, such as translation speeds, of *E. coli* are available in the future.

We remark that in the absence of heterogeneous translation speeds and protein degradation, the mass fraction of protein *i, ϕ_i_* must equal the ribosome allocation *χ_i_*. Indeed, these two datasets are often highly correlated among proteins in *E. coli* [12, 25]. However, in our more realistic models, *ϕ_i_* depends on the translation speed and protein degradation rate. Given the same *χ_i_*, proteins with higher translation speeds or lower degradation rates should have higher mass fractions (Appendix A). We note that using the current genome-wide datasets of *S. cerevisiae*, the predicted protein mass fractions *ϕ_i,pre_* based on the ribosome allocations *χ_i_* [14], the translation speeds *k_i_* [9], and the protein degradation rates *α_i_* [11] do not correlate strong enough with the measured *ϕ_i_* as expected. We note that these datasets are from different references, and the deviation is likely due to the noise in the measurements of *k_i_* (Supplementary Table S2). We expect our theories to be further verified when more accurate measurements of translation speeds are available.

For simplicity, in this work, we assume that the translation speeds and protein degradation rates are invariant as the nutrient quality changes. Therefore, we can use the two correlation coefficients *I_χ,k_* and *I_ϕ,α_* to characterize a particular environment. We remark that our model can be generalized to more complex scenarios in which the translation speeds or protein degradation rates depend on the growth rate [7]. In this case, one just needs to include four additional environmental-specific parameters: *k_R_*, 〈*k*〉, *ρ_R_*, and 〈*α*〉.

## APPENDIX

### A. Derivation of Equation (2)

Based on the definition of *ϕ_i_*, the changing rates of *ϕ_i_* is

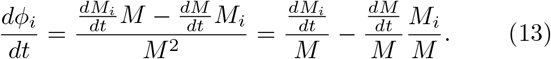

In the steady state, *ϕ_i_* doesn’t change so that Eq. (13) equals 0. Combined with the definition of growth rate and Eq. (1), we obtain

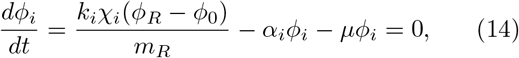

which leads to Eq. (2). In the steady state, we can write *ϕ_i_* using Eq. (2) as

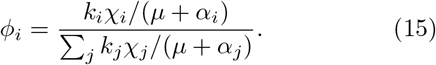

We can also rewrite Eq. (2) using Σ_*i*_ *ϕ_i_* = 1 as

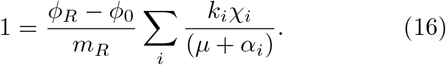

### B. Derivation of Equation (3)

We rewrite Eq. (2) as

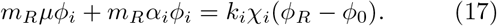

We then sum up for all proteins and obtain

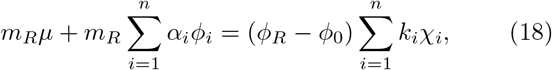

which leads to Eq. (3).

### C. Derivation of Equation (4)

In deriving Eq. (4), we neglect protein degradation and rewrite Eq. (3) as

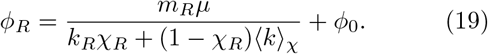

Meanwhile, we compute the growth rate using the autocatalytic nature of ribosomal proteins,

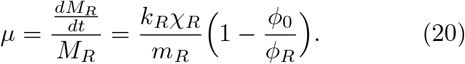

The above equation allows us to replace *χ_R_* by *μ* in Eq. (19), from which we obtain Eq. (4).

### D. Derivation of the full model

In this section we derive the full model considering both the heterogeneities in the translation speeds and protein degradation rates. We rewrite Eq. (3) in the main text as

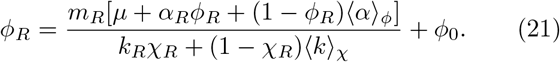

Meanwhile, the growth rate is

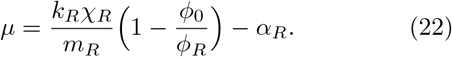

Combining Eq. (21) and Eq. (22) allows us to solve *ϕ_R_* as a function of *μ* and we obtain Eq. (12)

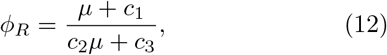

where

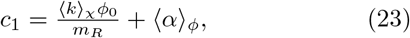

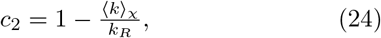

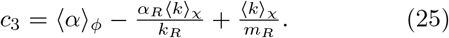

It is straightforward to find that the condition for Eq. (12) to be monotonically increasing is that *c*_3_ > *c*_1_*c*_2_. Using the above expressions, we find that

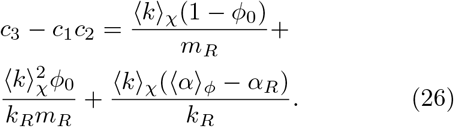

We find that the first two terms are always positive, and the last term is positive as long as *I_α,ϕ_* is not too close to −1. Therefore, the *ϕ_R_*(*μ*) curve must be monotonically increasing. It is straightforward to find that the second derivative of the *ϕ_R_*(*μ*) curve is proportional to (*c*_1_*c*_2_ – *c*_3_)*c*_2_, which is always positive as long as *I_χ,k_* is not too close to −1.

### E. Details of the numerical simulations

We summarize the parameters we use in the numerical simulations in Supplementary Table S1. We consider a cell with 4000 genes. We set the elongation speed *k_i_* and the degradation rates *α_i_* of non-ribosomal genes to follow lognormal distributions. We set *k_R_* = 2.07 × 10^4^ Da/min, 〈*k*〉 = 4.80 × 10^4^ Da/min, *α_R_* = 4.83 × 10^−4^ min^-1^, and 〈*α*〉 = 1.10 × 10^-3^ min^-1^ as the experimentally measured values of *S. cerevisiae* [9, 11]. The coefficients of variation (CV) of the lognormal distributions can be found in Supplementary Table S1. In all simulations, we set *ϕ*_0_ = 0.08. We note that in Figure 2a, we set *α_i_* = 0 for all proteins and in Figure 2b, we set *k_i_* = 〈*k*〉 for all proteins. We note that for given *I_χ,k_* and *I_ϕ,α_*, *k_i_* and *α_i_* are fixed for environments with different *χ_i_*.

To simulate a random environment, we generate a random *χ_R_*. Meanwhile, a lognormal distribution of *χ_i_* of non-ribosomal genes is also randomly generated. The CV of the lognormal distribution is included in Supplementary Table S1. We then search for the *ϕ_R_* and *μ* that simultaneously satisfy Eq. (22) and Eq. (16). *ϕ_i_*, *I_χ,k_* and *I_ϕ,α_* are then calculated using Eq. (3), Eq. (7) and Eq. (11), respectively. For a chosen pair of *I_χ,k_* and *I_ϕ,α_*, the predicted *ϕ_R_*(*μ*) curve is obtained using Eq. (12).

To obtain Figure. 2d, g, we randomly sample 20 points from Fig. 2c, f respectively, fit them using Eq. (12), and calculate the resulting RMSE. We repeat the above process 5000 times.

### F. Details of the experimental data analysis

For the ribosome profiling data [14], we first trim the adapter with Cutadapt (version 3.4) [26]. Then we use Bowtie2 (version 2.4.2) [27] to eliminate ribosomal RNAs (rRNA) as mentioned in [28]. The cleaned reads are then mapped to *S. cerevisiae* genome R64.1.1 with HISAT2 (version 2.2.1) [29]. Read counts are then generated with featureCount (version 2.0.1) [30]. The ribosome allocation *χ_i_* is calculated based on the count fraction.

For the proteomics data [14], we perform the absolute quantification (or the in-sample relative quantification) of proteins based on the intensities of peptides using xTop (version 1.2) [12]. The intensity ratio of 2 proteins in the same sample of proteomics data does not directly represent the real abundance (either the mass or the copy number) ratio so that the abundance fraction can not be replaced with the intensity fraction [12, 31]. XTop is a novel software that accurately calculates the in-sample relative protein copy number with the maximum a posteriori probability (MAP) algorithm [12]. We then calculate all proteins’ mass fraction *ϕ_i_* with the xTop results and the protein molecular mass. In [12], the authors further calibrated *ϕ_i_* with ribosome profiling data assuming homogeneous *k_i_*. In this work, we alternatively calibrate *ϕ_i_* with *L*^-0.57^ where *L* is the protein length, as mentioned in [12]. Calibration with *L*^-0.57^ is independent of ribosome profiling data, although it reduces the distance between *χ_i_* and calibrated *ϕ_i_* [12]. We also show the result with calibration of *L*^-1^ or without calibration in Supplementary Figure S1b, c. To compute *ϕ_R_*, we sum up the *ϕ_i_* of all proteins annotated as the cytoplasmic ribosomal protein in Saccharomyces Genome Database (SGD).

For the elongation speed *k_i_*, we first calculate *v_i_* as mentioned in [9]. *k_i_* is then calculated using the relationship *k_i_* = *v_i_a_i_*. For the degradation rate *α_i_*, data is obtained from [11]. We calculate the experimental *I_χ,k_*, *I_ϕ,α_*, 〈*k*〉 and 〈*α*〉 for non-ribosomal genes that exist in all data sets of *χ_i_, ϕ_i_, k_i_* and *α_i_*. We also calculate the *χ*-averaged *k* of ribosomal proteins as *k_R_* and *ϕ*-averaged, *α* of ribosomal proteins as *α_R_*.

For the molecular mass of the ribosome, we calculate the effective *m_R_*. Considering the efficiency of the mass spectrometry (MS), not all proteins can be detected. Therefore, we define the effective *m_R_* as the molecular weights of ribosomal proteins detected in the proteome. Because most of the ribosomal proteins can be expressed by two paralogous genes in *S. cerevisiae*, we count the average molecular mass when both proteins of the paralogs are detected in the proteome. We also show our predictions of *ϕ_R_* using the real ribosome mass (*m_R_* = 1.40e6 Da) in Supplementary Figure S1a.

For the growth rate μ, it is obtained from the growth curve, OD_600_ versus time with the method mentioned in [32]. Briefly, the slopes of ln(OD_600_) versus time in 5-point windows are calculated. Then windows with slopes that are at least 95% of the maximum slope are extracted. The slope of points within these windows is calculated as the growth rate. With these results, we predict the corresponding *ϕ_R_*(*μ*) curves and compare them with the experimental data points.

We further calculate the predicted mass fraction *ϕ_i,pre_* of non-ribosomal proteins with Eq. (15). Pearson correlation coefficients *ρ* between *ϕ_i,pre_* and *ϕ_i_* are calculated. We also compute *ρ* under the assumptions that *α_i_* =0 > or *k_i_* = 〈*k*〉 (Supplementary Table S2).

For GSEA analysis, we first perform the differential expression analysis on the ribosome profiling data of WT or i ΔNaa10 cells using the package DEseq2 (version 1.24.0) [33] in R (version 3.6.1). The log_2_ fold changes of counts when cells changed from SC+2% glucose to SC+2% glycerol as well as the FDR q values are calculated. Ribosomal genes and genes with FDR q value > 0.05 are eliminated. We then pick out genes that also exist in the data sets of *k_i_*. GSEA on these genes are then performed twice using the R package clusterProfiler (version 3.12.0) [34] and org.Sc.sgd.db (version 3.8.2) [35]. In the first GSEA, genes are ordered by the log_2_ fold change (denoted as log_2_FC-ordered GSEA). In the second GSEA, genes are ordered by *k_i_* (denoted as *k_i_*-ordered GSEA). We then find the common gene sets from GO database [17, 18] enriched in these two GSEA. The cut-off criteria are set as the p value < 0.05 and the FDR q value < 0.25. The number of permutations used in the analysis is 1e5.

### G. Details of fitting in Figure 5

Nonlinear fitting is performed with MATLAB (version R2020b). We obtain the fitting parameters *c*_1_, *c*_2_ and *c*_3_ with their 95% confidence intervals, and then compute *ϕ*_0_, *k_R_* and 〈*k*〉_*χ*_ using Eqs. (23, 24, 25). To compute the ranges of these values, we numerically find the maximum and the minimum value of the multivariate functions *ϕ*_0_(*c*_1_, *c*_2_, *c*_3_), *k_R_*(*c*_1_,*c*_2_,*c*_3_) and 〈*k*〉_*χ*_(*c*_1_, *c*_2_, *c*_3_) as their upper and lower bounds, where the ranges of *c*_1_, *c*_2_ and *c*_3_ are their 95% confidence intervals.

### H. A summary of the variables used in this work

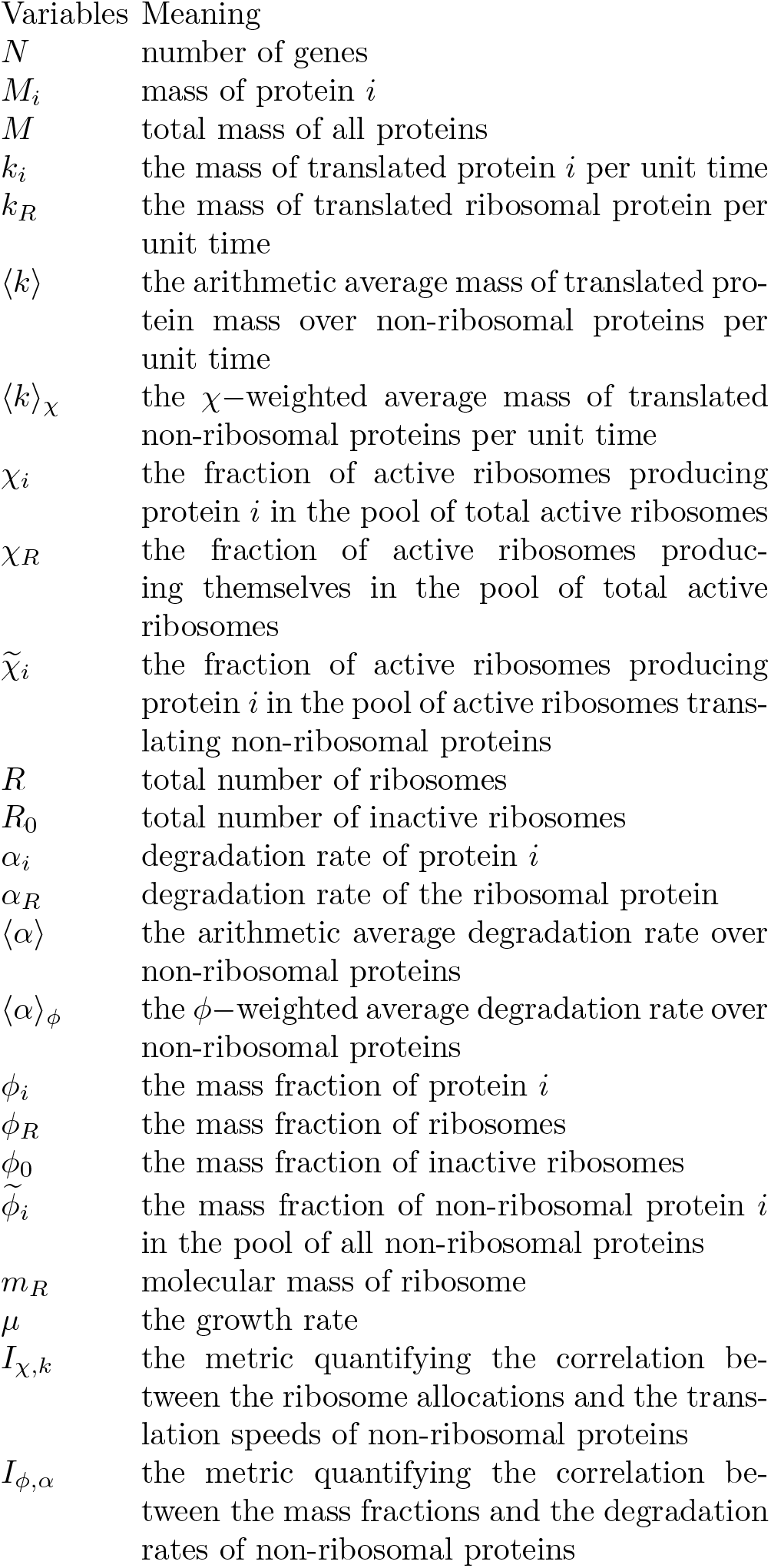

## Notes

### Competing Interest Statement

The authors have declared no competing interest.

## References

[1] F. C. Neidhardt and B. Magasanik, Studies on the role of ribonucleic acid in the growth of bacteria, Biochimica et biophysica acta 42, 99 (1960).

[2] O. Maaløe, Regulation of the protein-synthesizing machinery—ribosomes, trna, factors, and so on, in Biological regulation and development (Springer, 1979) pp. 487–542.

[3] H. Bremer and P. P. Dennis, Modulation of chemical composition and other parameters of the cell at different exponential growth rates, EcoSal Plus 3 (2008).

[4] M. Scott, C. W. Gunderson, E. M. Mateescu, Z. Zhang, and T. Hwa, Interdependence of Cell Growth and Gene Expression: Origins and Consequences, Science 330, 1099 (2010).

[5] S. Hui, J. M. Silverman, S. S. Chen, D. W. Erickson, M. Basan, J. Wang, T. Hwa, and J. R. Williamson, Quantitative proteomic analysis reveals a simple strategy of global resource allocation in bacteria, Molecular Systems Biology 11, 784 (2015).

[6] E. Metzl-Raz, M. Kafri, G. Yaakov, I. Soifer, Y. Gurvich, and N. Barkai, Principles of cellular resource allocation revealed by condition-dependent proteome profiling, Elife 6, e28034 (2017).

[7] X. Dai, M. Zhu, M. Warren, R. Balakrishnan, V. Patsalo, H. Okano, J. R. Williamson, K. Fredrick, Y.-P. Wang, and T. Hwa, Reduction of translating ribosomes enables escherichia coli to maintain elongation rates during slow growth, Nature microbiology 2, 1 (2016).

[8] S. Klumpp, J. Dong, and T. Hwa, On ribosome load, codon bias and protein abundance, PLoS ONE 7, e48542 (2012).

[9] A. Riba, N. Di Nanni, N. Mittal, E. Arhné, A. Schmidt, and M. Zavolan, Protein synthesis rates and ribosome occupancies reveal determinants of translation elongation rates, Proceedings of the national academy of sciences 116, 15023 (2019).

[10] O. Gefen, O. Fridman, I. Ronin, and N. Q. Balaban, Direct observation of single stationary-phase bacteria reveals a surprisingly long period of constant protein production activity, Proceedings of the National Academy of Sciences 111, 556 (2014).

[11] P.-J. Lahtvee, B. J. Sánchez, A. Smialowska, S. Kasvandik, I. E. Elsemman, F. Gatto, and J. Nielsen, Absolute quantification of protein and mrna abundances demonstrate variability in gene-specific translation efficiency in yeast, Cell Systems 4, 495 (2017).

[12] M. Mori, Z. Zhang, A. Banaei-Esfahani, J.-B. Lalanne, H. Okano, B. C. Collins, A. Schmidt, O. T. Schubert, D.-S. Lee, G.-W. Li, R. Aebersold, T. Hwa, and C. Ludwig, From coarse to fine: the absolute escherichia coli proteome under diverse growth conditions, Molecular Systems Biology 17, e9536 (2021).

[13] G.-W. Li, D. Burkhardt, C. Gross, and J. S. Weissman, Quantifying absolute protein synthesis rates reveals principles underlying allocation of cellular resources, Cell 157, 624 (2014).

[14] U. A. Friedrich, M. Zedan, B. Hessling, K. Fenzl, L. Gillet, J. Barry, M. Knop, G. Kramer, and B. Bukau, N*α*-terminal acetylation of proteins by nata and natb serves distinct physiological roles in saccharomyces cerevisiae, Cell Reports 34, 108711 (2021).

[15] A. Subramanian, P. Tamayo, V. K. Mootha, S. Mukherjee, B. L. Ebert, M. A. Gillette, A. Paulovich, S. L. Pomeroy, T. R. Golub, E. S. Lander, and J. P. Mesirov, Gene set enrichment analysis: A knowledge-based approach for interpreting genome-wide expression profiles, Proceedings of the National Academy of Sciences 102, 15545 (2005).

[16] V. K. Mootha, C. M. Lindgren, K.-F. Eriksson, A. Subramanian, S. Sihag, J. Lehar, P. Puigserver, E. Carlsson, M. Ridderstrále, E. Laurila, and et al., Pgc-1alpha-responsive genes involved in oxidative phosphorylation are coordinately downregulated in human diabetes, Nature Genetics 34, 267–273 (2003).

[17] M. Ashburner, C. A. Ball, J. A. Blake, D. Botstein, H. Butler, J. M. Cherry, A. P. Davis, K. Dolinski, S. S. Dwight, J. T. Eppig, M. A. Harris, D. P. Hill, L. Issel-Tarver, A. Kasarskis, S. Lewis, J. C. Matese, J. E. Richardson, M. Ringwald, G. M. Rubin, and G. Sherlock, Gene ontology: tool for the unification of biology, Nature Genetics 25, 25 (2000).

[18] S. Carbon, E. Douglass, B. M. Good, D. R. Unni, N. L. Harris, C. J. Mungall, S. Basu, R. L. Chisholm, R. J. Dodson, E. Hartline, P. Fey, P. D. Thomas, L.-P. Albou, D. Ebert, M. J. Kesling, H. Mi, A. Muruganujan, X. Huang, T. Mushayahama, S. A. LaBonte, D. A. Siegele, G. Antonazzo, H. Attrill, N. H. Brown, P. Garapati, S. J. Marygold, V. Trovisco, G. dos Santos, K. Falls, C. Tabone, P. Zhou, J. L. Goodman, V. B. Strelets, J. Thurmond, P. Garmiri, R. Ishtiaq, M. Rodríguez-Lopez, M. L. Acencio, M. Kuiper, A. Lægreid, C. Logie, R. C. Lovering, B. Kramarz, S. C. C. Saverimuttu, S. M. Pinheiro, H. Gunn, R. Su, K. E. Thurlow, M. Chibucos, M. Giglio, S. Nadendla, J. Munro, R. Jackson, M. J. Duesbury, N. Del-Toro, B. H. M. Meldal, K. Paneerselvam, L. Perfetto, P. Porras, S. Orchard, A. Shrivastava, H.-Y. Chang, R. D. Finn, A. L. Mitchell, N. D. Rawlings, L. Richardson, A. Sangrador-Vegas, J. A. Blake, K. R. Christie, M. E. Dolan, H. J. Drabkin, D. P. Hill, L. Ni, D. M. Sitnikov, M. A. Harris, S. G. Oliver, K. Rutherford, V. Wood, J. Hayles, J. Bähler, E. R. Bolton, J. L. D. Pons, M. R. Dwinell, G. T. Hayman, M. L. Kaldunski, A. E. Kwitek, S. J. F. Laulederkind, C. Plasterer, M. A. Tutaj, M. Vedi, S.-J. Wang, P. D’Eustachio, L. Matthews, J. P. Balhoff, S. A. Aleksander, M. J. Alexander, J. M. Cherry, S. R. Engel, F. Gondwe, K. Karra, S. R. Miyasato, R. S. Nash, M. Simison, M. S. Skrzypek, S. Weng, E. D. Wong, M. Feuermann, P. Gaudet, A. Morgat, E. Bakker, T. Z. Berardini, L. Reiser, S. Subramaniam, E. Huala, C. N. Arighi, A. Auchincloss, K. Axelsen, G. Argoud-Puy, A. Bateman, M.-C. Blatter, E. Boutet, E. Bowler, L. Breuza, A. Bridge, R. Britto, H. Bye-A-Jee, C. C. Casas, E. Coudert, P. Denny, A. Estreicher, M. L. Famiglietti, G. Georghiou, A. Gos, N. Gruaz-Gumowski, E. Hatton-Ellis, C. Hulo, A. Ignatchenko, F. Jungo, K. Laiho, P. L. Mercier, D. Lieberherr, A. Lock, Y. Lussi, A. MacDougall, M. Magrane, M. J. Martin, P. Masson, D. A. Natale, N. Hyka-Nouspikel, S. Orchard, I. Pedruzzi, L. Pourcel, S. Poux, S. Pundir, C. Rivoire, E. Speretta, S. Sundaram, N. Tyagi, K. Warner, R. Zaru, C. H. Wu, A. D. Diehl, J. N. Chan, C. Grove, R. Y. N. Lee, H.-M. Muller, D. Raciti, K. V. Auken, P. W. Sternberg, M. Berriman, M. Paulini, K. Howe, S. Gao, A. Wright, L. Stein, D. G. Howe, S. Toro, M. Westerfield, P. Jaiswal, L. Cooper, and J. Elser, The gene ontology resource: enriching a GOld mine, Nucleic Acids Research 49, D325 (2020).

[19] J. Gutin, A. Sadeh, A. Rahat, A. Aharoni, and N. Friedman, Condition-specific genetic interaction maps reveal crosstalk between the camp/pka and the hog mapk pathways in the activation of the general stress response, Molecular Systems Biology 11, 829 (2015).

[20] D. W. Erickson, S. J. Schink, V. Patsalo, J. R. Williamson, U. Gerland, and T. Hwa, A global resource allocation strategy governs growth transition kinetics of escherichia coli, 551, 119 (2017).

[21] A. L. Goldberg and A. C. S. John, Intracellular protein degradation in mammalian and bacterial cells: Part 2, 45, 747 (1976).

[22] A. Arkin, J. Ross, and H. H. McAdams, Stochastic kinetic analysis of developmental pathway bifurcation in phage lambda-infected escherichia coli cells, 149, 1633 (1998).

[23] M. Gouy and R. Grantham, Polypeptide elongation and trna cycling in escherichia coli: a dynamic approach, FEBS letters 115, 151 (1980).

[24] T. V. Karpinets, D. J. Greenwood, C. E. Sams, and J. T. Ammons, RNA:protein ratio of the unicellular organism as a characteristic of phosphorous and nitrogen stoichiometry and of the cellular requirement of ribosomes for protein synthesis 4, 10.1186/1741-7007-4-30 (2006).

[25] T.-Y. Liu, H. H. Huang, D. Wheeler, Y. Xu, J. A. Wells, Y. S. Song, and A. P. Wiita, Time-resolved proteomics extends ribosome profiling-based measurements of protein synthesis dynamics, Cell Systems 4, 636 (2017).

[26] M. Martin, Cutadapt removes adapter sequences from high-throughput sequencing reads, EMBnet.journal 17, 10 (2011).

[27] B. Langmead and S. L. Salzberg, Fast gapped-read alignment with bowtie 2, Nature methods 9, 357 (2012).

[28] C. V. Galmozzi, D. Merker, U. A. Friedrich, K. Döring, and G. Kramer, Selective ribosome profiling to study interactions of translating ribosomes in yeast, Nature protocols 14, 2279 (2019).

[29] D. Kim, J. M. Paggi, C. Park, C. Bennett, and S. L. Salzberg, Graph-based genome alignment and genotyping with hisat2 and hisat-genotype, Nature biotechnology 37, 907 (2019).

[30] Y. Liao, G. K. Smyth, and W. Shi, featureCounts: an efficient general purpose program for assigning sequence reads to genomic features, Bioinformatics 30, 923 (2013).

[31] F. Calderón-Celis, J. R. Encinar, and A. Sanz-Medel, Standardization approaches in absolute quantitative proteomics with mass spectrometry, Mass spectrometry reviews 37, 715 (2018).

[32] B. G. Hall, H. Acar, A. Nandipati, and M. Barlow, Growth Rates Made Easy, Molecular Biology and Evolution 31, 232 (2013).

[33] M. I. Love, W. Huber, and S. Anders, Moderated estimation of fold change and dispersion for rna-seq data with deseq2, Genome Biology 15, 550 (2014).

[34] G. Yu, L.-G. Wang, Y. Han, and Q.-Y. He, clusterProfiler: an r package for comparing biological themes among gene clusters, OMICS: A Journal of Integrative Biology 16, 284 (2012).

[35] M. Carlson, org.sc.sgd.db: Genome wide annotation for yeast, (2019), r package version 3.8.2.

