## Supplementary material for "Universality and non-universality of the growth law": SI_6.pdf

Table S1: A summary of the parameters used in the numerical simulations if not mentioned in the text.

| Parameters | Meaning | Values |
|---|---|---|
| $N$ | number of genes | 4.00e3 |
| $m_R$ | molecular mass of the ribosome | 1.40e6 Da |
| $\phi_0$ | the mass fraction of inactive ribosomes | 8.00e-2 |
| $\langle k \rangle$ | the average mass of translated non-ribosomal proteins per unit time | 4.80e4 Da/min |
| $CV_k$ | The coefficient of variation for the distribution of $k_i$ | 1.97 |
| $k_R$ | the mass of translated ribosomal protein per unit time | 2.07e4 Da/min |
| $\langle \alpha \rangle$ | the average degradation rate of non-ribosomal proteins | 1.10e-3 1/min |
| $CV_\alpha$ | The coefficient of variation for the distribution of $alpha_i$ | 1.76 |
| $\alpha_R$ | degradation rate of the ribosomal protein | 4.83e-4 1/min |
| $CV_\chi$ | The coefficient of variation for the distribution of $\chi_i$ | 4.5-5.5 |

Table S2: A summary of the Pearson correlations between the predicted $\phi_i$ and experimental $\phi_i$. We also show the correlation between the experimental $\chi_i$ and $\phi_i$. The calculation of the predicted $\phi_i$ can be found in the Appendix F of the main text.

| Data | WT+2%glu | WT+2%gly | ΔNaa10+2%glu | ΔNaa10+2%gly |
|---|---|---|---|---|
| $\rho_{\phi,\chi}$ | 0.82 | 0.75 | 0.83 | 0.70 |
| $\rho_{\phi_{pre},\phi}$ | 0.68 | 0.26 | 0.79 | 0.23 |
| $\rho_{\phi_{pre},\phi}$ with $\alpha_i = 0$ | 0.69 | 0.23 | 0.80 | 0.22 |
| $\rho_{\phi_{pre},\phi}$ with $k_i = \langle k \rangle$ | 0.82 | 0.76 | 0.82 | 0.74 |
| $\rho_{\phi_{pre},\phi}$ with $\alpha_i = 0$ and $k_i = \langle k \rangle$ | 0.82 | 0.75 | 0.83 | 0.70 |

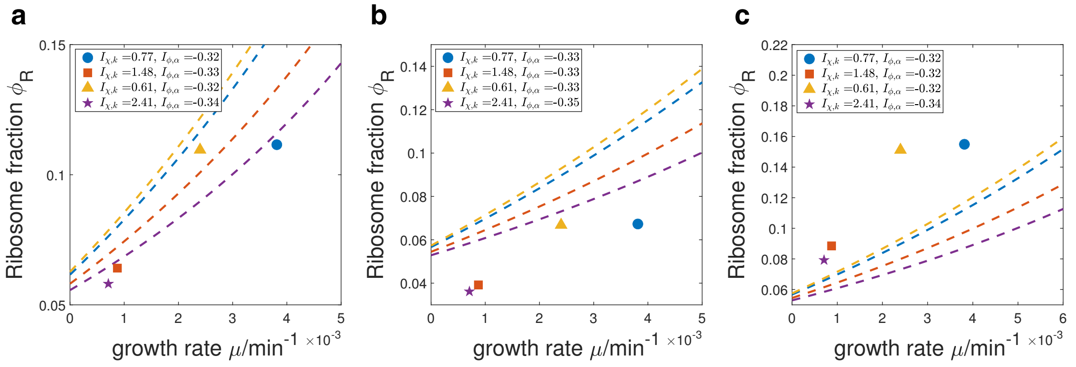


Figure S1: Experimental tests of the theoretical predictions of $\phi_R$ with different data processing procedures. $\phi_0 = 4.80e-2$ are used in all the cases. (a) The case in which the actual ribosomal mass $m_R = 1.40e6$ Da is used in the predictions. (b) The case in which the experimental $\phi_i$ is not calibrated. (c) The case in which the experimental $\phi_i$ is calibrated with $L^{-1}$.

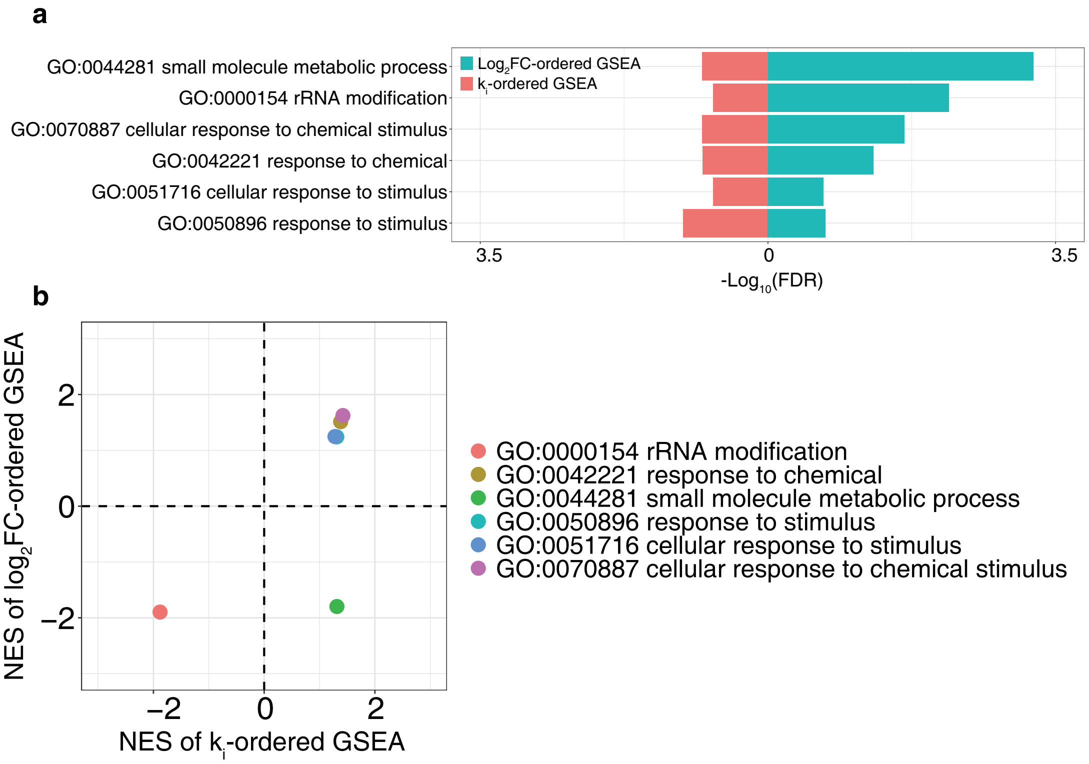


Figure S2: GSEA results of ΔNaa10 cells. (a) The enriched gene sets with their false discovery rate (FDR) q values. (b) The normalized enrichment score (NES) of GSEA of enriched gene sets. A positive NES of $k_i$-ordered GSEA means that the genes in the corresponding gene set are enriched in the regime of higher $k_i$. A positive NES of $\log_2$FC-ordered GSEA means that the genes in the corresponding gene set are enriched in the regime of increasing $\chi_i$ when the nutrient changes from glucose to glycerol.

[1] Eyal Metzl-Raz, Moshe Kafri, Gilad Yaakov, Ilya Soifer, Yonat Gurvich, and Naama Barkai, "Principles of cellular resource allocation revealed by condition-dependent proteome profiling," Elife **6**, e28034 (2017).

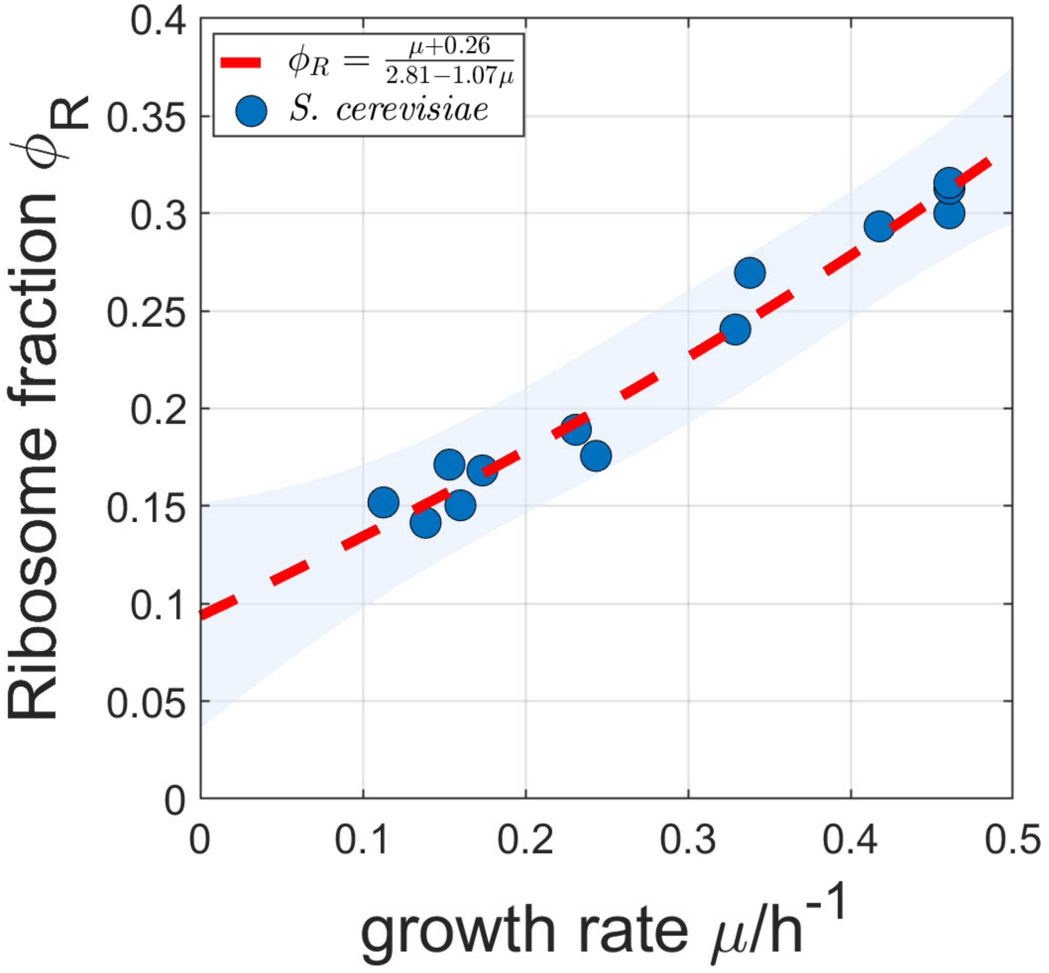


Figure S3: The non-linear fitting of the C-limiting data from Ref. [1].
